# A Variational Principle for the Emergence of Neural Geometry Underlying Bayesian-Like Behavior

**DOI:** 10.64898/2026.11.11.765947

**Authors:** Luis Serrano-Fernández, Néstor Parga

## Abstract

Neural population geometry constrains how task variables are represented and read out into behavior. We propose a variational principle in which learning balances task accuracy and representational smoothness under stimulus-dependent weighting. This balance links stimulus statistics to the shape of a latent neural coordinate. For magnitude estimation, the principle predicts distortions near the edges of bounded ensembles, global affine contraction for Gaussian ensembles, and warping toward high-probability regions for bimodal ensembles. We test the bounded-uniform case in recurrent neural networks trained to reproduce time intervals. Their learned geometries occupy three regimes as the boundary distortions change from too narrow to resolve, to measurable, to extending across the stimulus range. A shape parameter selected from an empirical neural coordinate predicts the nonlinear component of mean behavior through an affine readout, with only behavioral gain and offset fitted afterward. In this bounded-uniform setting, the variational coordinate and Bayesian least-squares mean response have closely matching shapes after affine normalization. These results suggest a variational account of how stimulus statistics shape neural geometry and yield Bayesian-like behavior.

## Introduction

In many cognitive tasks, neural populations encode stimuli, choices, memories, and motor plans in low-dimensional manifolds embedded in high-dimensional state space [1]. Downstream computations read out coordinates on these manifolds, making the relation between learned geometry and behavior a central problem in theoretical and experimental systems neuroscience [2–5].

We propose a variational principle that determines a latent coordinate of the learned stimulus manifold by balancing task accuracy and smoothness under stimulus-dependent weighting. We develop the framework for magnitude-reconstruction tasks [4, 6, 7], which allow the behavioral consequences of geometric distortions to be analyzed explicitly. Finally we compare the variational model with the Bayesian model [7].

### State-space geometry and the stimulus manifold

For a stimulus value *s*, population activity follows a trajectory **x**(*t*; *s*) in *N* -dimensional state space. We compare stimuli at a common computational stage, indexed by an encoding phase *φ*, and define

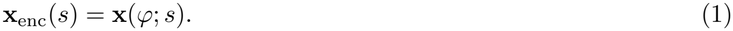

These states trace the measured stimulus manifold *ℳ*_enc_ = *{***x**_enc_(*s*) : *s ∈ S}* (Fig. 1(a)). To describe stimulus encoding along this manifold, we introduce a scalar latent coordinate *u*(*s*), determined by the variational principle below. A nonlinear map *s ↦ u*(*s*) assigns unequal latent increments to equally spaced stimuli (Fig. 1(b)). The predicted mean behavioral response is

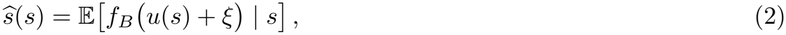

where *f*_*B*_ is the downstream readout and *ξ* denotes noise.

**FIG. 1.**
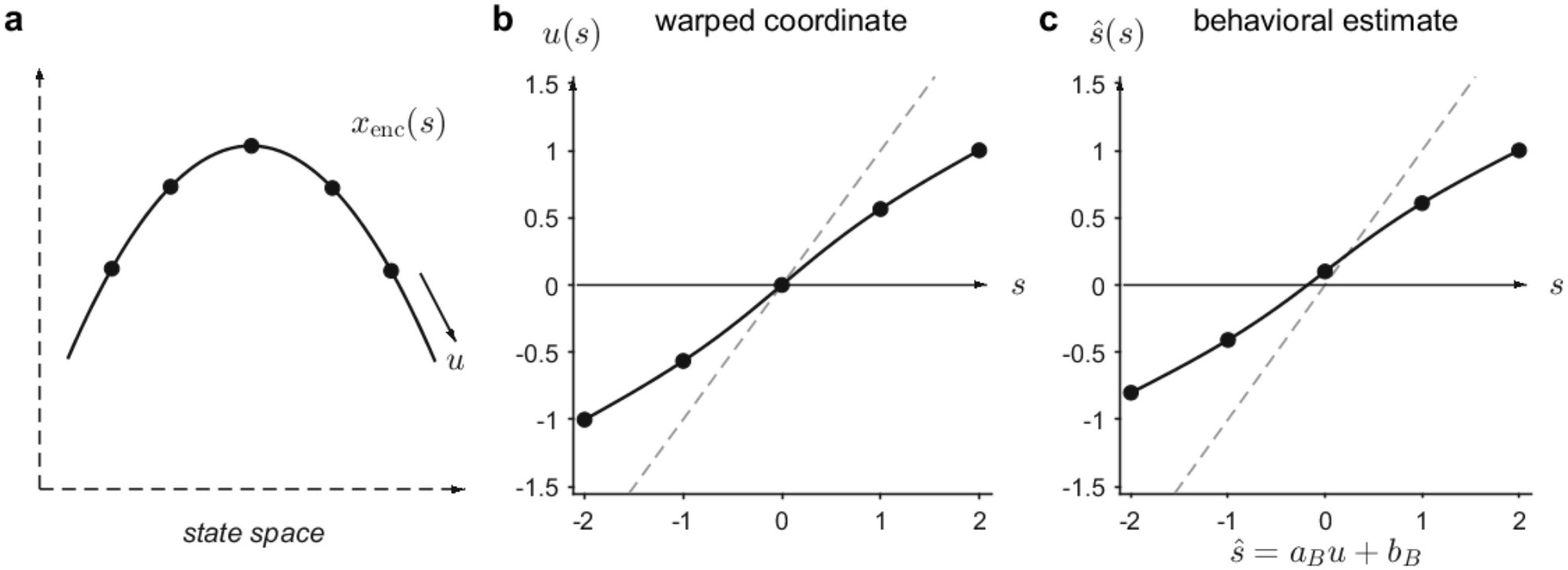
Neural geometry and behavioral readout. (a) Population states **x**_enc_(*s*) at a common computational phase *ϕ* define the stimulus manifold. (b) A nonlinear latent coordinate *u*(*s*) assigns unequal increments to equally spaced stimuli. (c) An affine readout 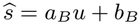 transfers this shape to behavior. Dashed lines in (b,c) denote identity maps.

### Variational principle

Let *L*_task_(*s, u*) denote the coarse-grained task loss. We balance this loss against a penalty on variation across nearby stimuli through the functional

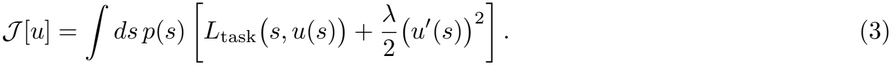

Here the normalized density *p*(*s*) weights each stimulus’s contribution during learning, and *λ* controls the smoothness penalty.

### Magnitude-estimation tasks

In magnitude reconstruction or local estimation, a stimulus magnitude *s*, such as a time interval, length, frequency, or intensity, must be estimated or reproduced from neural activity [4, 6, 7]. A robust phenomenon is contraction bias: small stimuli tend to be overestimated and large stimuli underestimated [8]. We ask whether this bias reflects representational geometry. The variational principle determines the latent coordinate *u*(*s*), whereas a downstream affine readout can modify its overall gain and offset. For a noiseless affine readout, 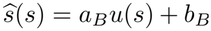, behavior inherits the latent shape up to gain and offset. We therefore refer to *Δ*(*s*) = *u*(*s*) *− s* as the geometric distortion and reserve behavioral bias for 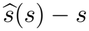.

For these tasks, we use a quadratic fidelity loss,

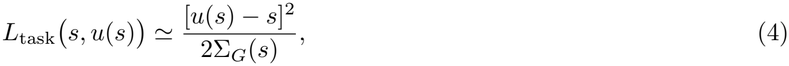

where Σ_*G*_(*s*) is the local inverse-fidelity scale: larger values weaken the anchoring of *u*(*s*) to the physical stimulus. The associated Euler–Lagrange equation is

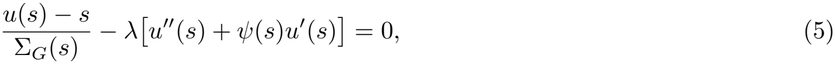

where *ψ*(*s*) = *∂*_*s*_ log *p*(*s*) is the score of the stimulus-weighting density.

#### Relating the variational coordinate to neural geometry

From the measured manifold *ℳ*_enc_, we extract an empirical scalar coordinate

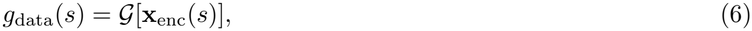

where *G* denotes a geometric analysis, such as projection onto an encoding axis or signed distance to a reference state or trajectory [9–12].

The fidelity loss anchors *u*(*s*) to the physical stimulus scale, whereas the origin, scale, and orientation of *g*_data_(*s*) depend on its construction. We therefore compare them through the affine observation model

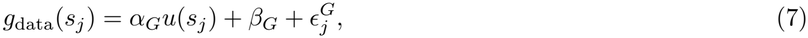

where 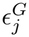 is the residual at stimulus *s*_*j*_.

### The weak-distortion regime

To obtain the leading departure from the identity map, we substitute the geometric distortion *Δ*(*s*) = *O*(*λ*) into Eq. (5) obtaining

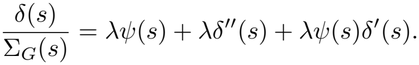

If *Δ* varies on a stimulus scale that remains finite and nonzero as *λ →* 0, then *Δ*^*′*^, *Δ*^*′′*^ = *O*(*λ*), and the last two terms are *O*(*λ*^2^). Thus, away from singular boundary layers,

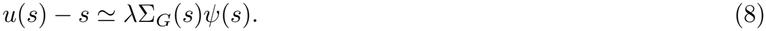

The leading distortion follows the score of *p*(*s*), weighted by the local inverse-fidelity scale, shifting the representation toward more strongly weighted stimuli. This regular interior approximation applies when *p*(*s*) varies smoothly on the scale of the distortion. Hard bounded support requires separate treatment.

#### 2. Gaussian weighting

Consider the distribution,

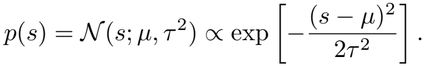

Then *τ* ^2^ *ψ*(*s*) = *−* (*s − µ*). For constant inverse fidelity Σ_*G*_(*s*) = Σ_0_, the Euler–Lagrange equation admits an exact global affine solution

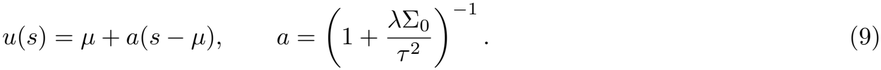

Thus 0 *< a <* 1. It contracts toward *µ* (Fig. 2(a)), but its gain can be absorbed by the affine observation model or behavioral readout. It therefore has no nonlinear signature distinguishable from an affine null.

**FIG. 2.**
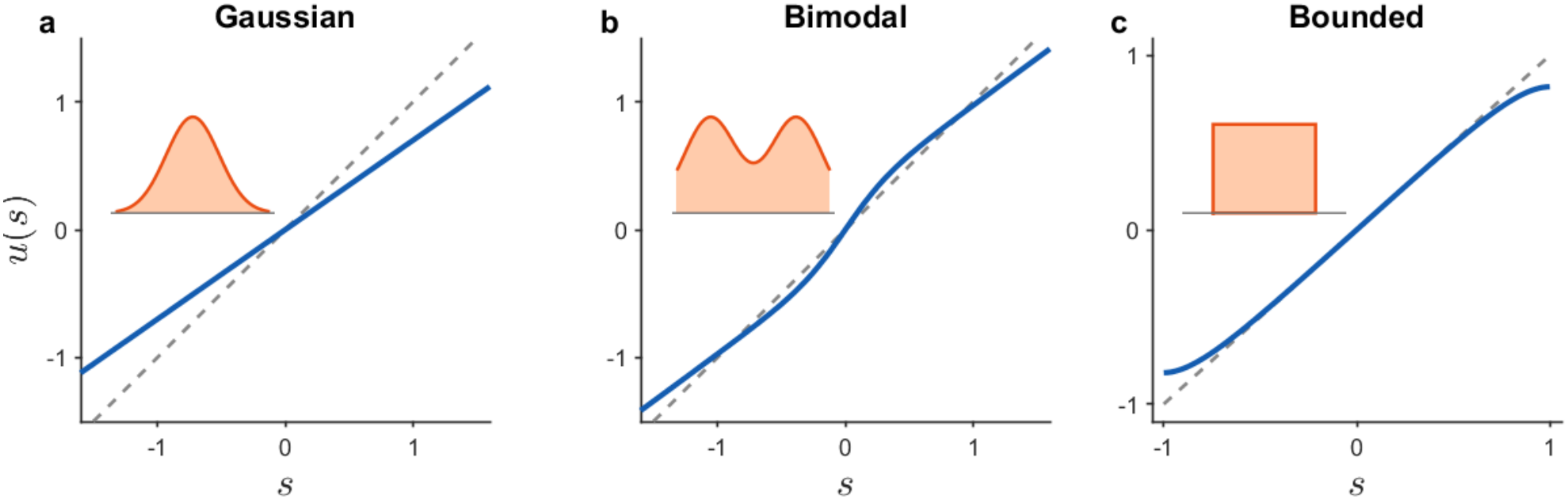
Variational solutions for different stimulus weightings. Blue curves show *u*(*s*), orange insets show *p*(*s*), and dashed lines denote *u* = *s*. For constant inverse fidelity, (a) Gaussian weighting gives affine contraction, (b) symmetric bimodal weighting produces warping toward the modes in the weak-distortion regime, and (c) bounded-uniform weighting produces separated boundary layers with a nearly linear interior. Dimensionless plotting parameters are *µ* = 0, *τ* = 0.55, and *a* = 0.70 in (a); *m* = 0.90, *τ* = 0.55, and *λ*Σ_0_ = 0.08 in (b); and *s*_*±*_ = *±*1 and *ℓ* = 0.18 in (c).

#### 2. Bimodal weighting

The same principle predicts more structured geometries when the stimulus ensemble is bimodal. For a symmetric bimodal distribution,

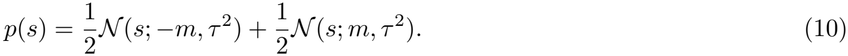

In the weak-distortion regime with constant inverse fidelity,

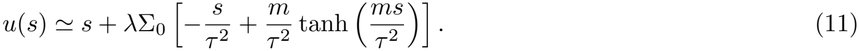

For *m > τ*, the center is a local minimum of *p*(*s*), and the leading distortion points toward the two modes (Fig. 2(b)). Indeed, near *s* = 0

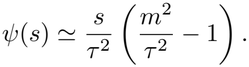

#### 3. Bounded-uniform weighting

Consider the stimulus-weighting density

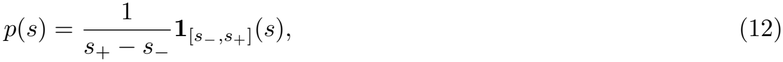

Its score vanishes in the interior of the support, *ψ*(*s*) = 0. Equation (8) therefore predicts *u*(*s*) *≃ s* in the bulk, but this regular approximation cannot satisfy the natural boundary conditions at the hard endpoints. The full Euler–Lagrange problem is instead

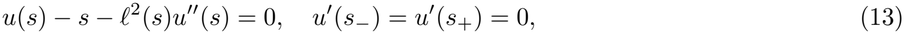

where *ℓ*^2^(*s*) = *λ*Σ_*G*_(*s*). For constant inverse fidelity the characteristic boundary-layer width is 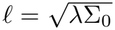. The exact solution is in the Supplemental Material; in the separated-boundary-layer regime, (*s*_+_ *− s*_*−*_)*/*(2*ℓ*) *≫* 1, it reduces to

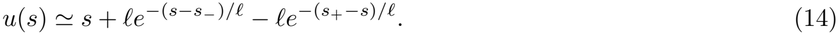

The distortions are localized near the endpoints, while the central region remains approximately linear with unit slope (Fig. 2(c)). When *ℓ* becomes comparable to the width of the stimulus range, however, the two boundary layers overlap and generate curvature throughout the interval.

### Trained recurrent neural networks (RNNs)

RNNs trained by supervised learning [13, 14] provide a natural dynamical setting for the assumptions underlying the variational principle with the task loss in Eq. (4). Near successful performance, a quadratic expansion of the output loss and a local linearization of the readout motivate the fidelity term. Smooth dependence of the coarse-grained population trajectories on stimulus magnitude motivates the regularity penalty.

### RNNs trained for time-interval reproduction

We next test the variational description in RNNs of spiking neurons trained on the time-interval reproduction task [7] using the algorithm for training spiking RNNs proposed in [13]. On each trial, a sample interval *s* is specified by two cues, indicating the beginning and end of the temporal interval, and must subsequently be reproduced. The networks are trained with a quadratic output loss on discrete intervals drawn from the bounded range [*s*_*−*_, *s*_+_]. We analyzed five networks at each training-ensemble center *C* = 650, 750, 850, 950 ms; C650-N05 denotes the fifth network at *C* = 650 ms. After training, their mean reproduction intervals, *s*_rep_(*s*) exhibit the usual contraction bias [7, 8]: short intervals are overproduced and long intervals are underproduced, with additional flattening near the limits of the trained range (symbols in Fig. 3 (d–f) for 3 selected networks and in Supplemental Fig. S1 for the 20 networks).

**FIG. 3.**
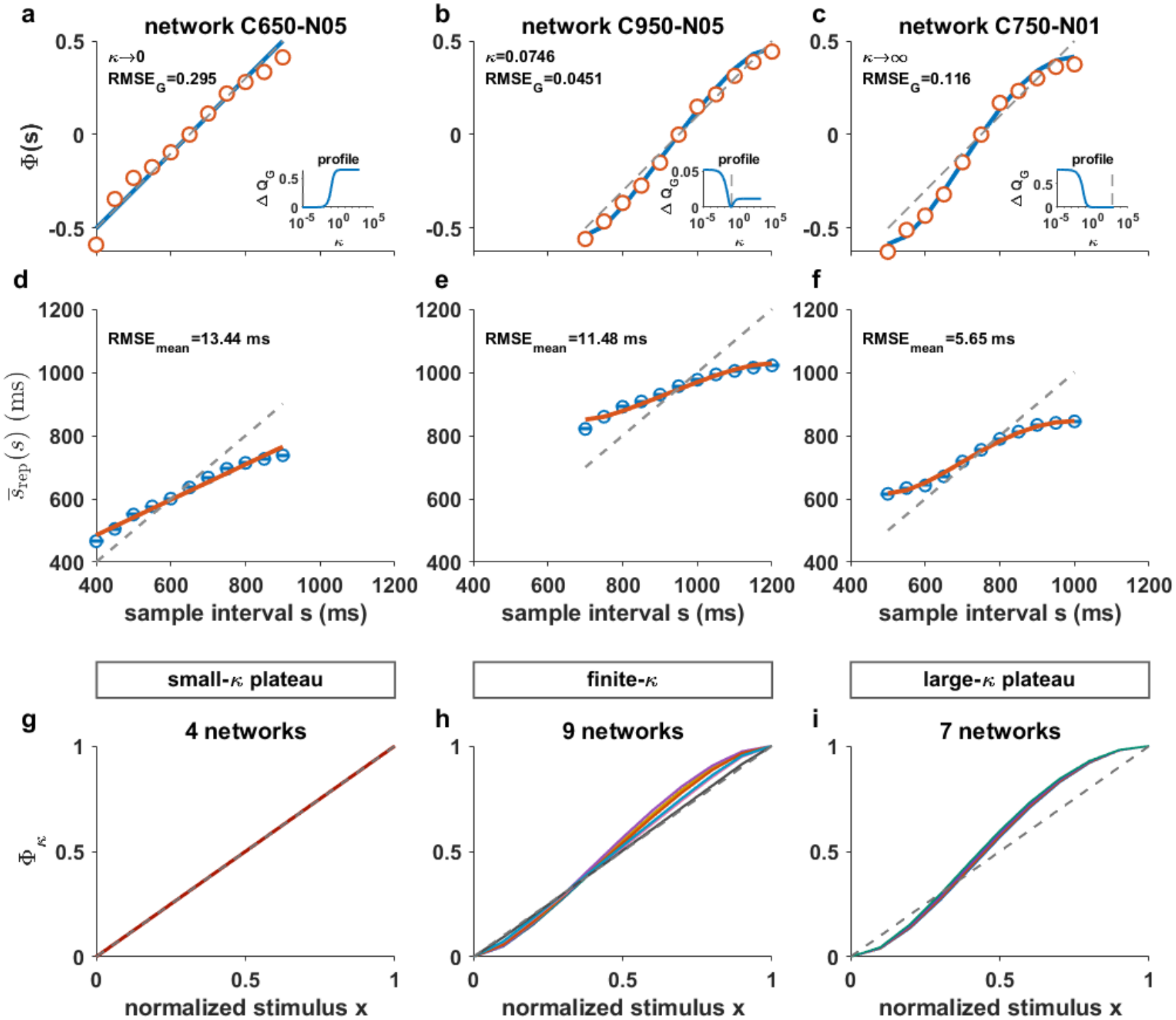
Neural geometry and behavior across three variational regimes. Columns represent small-, intermediate-, and large-*κ* regimes. (a–c) Normalized neural coordinates Φ_data_ (orange circles) and variational fits 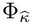 (blue). Insets show Δ*Q*_*G*_ = *Q*_*G*_ − min *Q*_*G*_ versus *κ* (logarithmic scale); plateaus indicate asymptotic regimes. (d–f) Mean reproduced intervals (blue circles; error bars, standard errors where available) and conditional predictions (orange), with 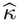 fixed by geometry and only behavioral gain and offset fitted. (g–i) Shapes for all 20 networks, grouped into the three regimes (4, 9, and 7 networks). Each curve represents one network, plotted as 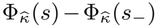 against *x* = (*s − s*_*−*_)*/*(*s*_+_ *− s*_*−*_). Dashed lines denote (*s − s*_*c*_)*/*(*s*_+_ *− s*_*−*_) in (a–c), veridical reproduction in (d–f), and *y* = *x* in (g–i). Annotations give 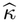 or its regime and RMSE_*G*_ in the original geometric coordinates (a–c), and RMSE across reproduction means in milliseconds (d–f).

For each sample interval *s*_*j*_, we evaluated the population trajectory at a common computational phase and constructed the empirical neural coordinate *g*_data_(*s*_*j*_) from signed distances to a reference trajectory corresponding to an interval *s*_*c*_ near the center of the trained range [5, 9, 10, 12] (Φ_data_(*s*) in Fig. 3 (a–c) for three representative networks, and the original *g*_data_(*s*) in Supplemental Fig. S2 for all 20 networks). Thus, each network provides two empirical curves: the neural geometric coordinate *g*_data_(*s*) and the behavioral response *s*_rep_(*s*). They will be compared with, respectively, the neural observation and behavioral readout of a common theoretical latent shape.

For this application, we take *p*(*s*) to be the continuous bounded-uniform approximation to the discrete training ensemble on [*s*_*−*_, *s*_+_]. Motivated by scalar timing [15–17], we assume 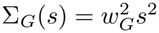 and define 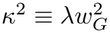.

For every value of *κ*, the variational coordinate *u*_*κ*_(*s*) is defined by the full boundary-value problem

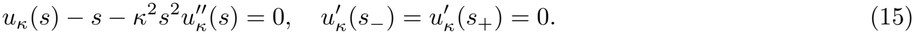

The notation *u*_*κ*_ makes explicit the dependence on *κ*. All variational fits use the exact solution of Eq. (15) (Supplemental Material). The local propagation length *ℓ*(*s*) = *κs* makes the upper boundary influence a broader range of intervals than the lower boundary.

Only the affine-invariant shape of *u*_*κ*_ can be compared with the empirical neural coordinate. Defining Δ*u*_*κ*_ *≡ u*_*κ*_(*s*_+_) *− u*_*κ*_(*s*_*−*_), we introduce

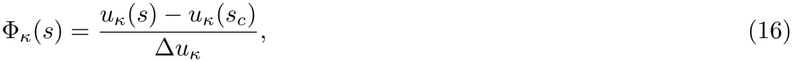

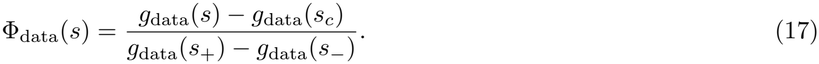

Φ_*κ*_ is invariant under arbitrary affine transformations of *u*_*κ*_ with nonzero gain. The neural observation model in Eq. (7) is equivalently written as

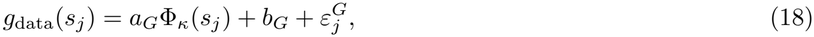

where *a*_*G*_ = *α*_*G*_Δ*u*_*κ*_ and *b*_*G*_ = *β*_*G*_ + *α*_*G*_*u*_*κ*_(*s*_*c*_).

For each network, we estimate *κ* from the neural geometric data by profiling out the neural gain and offset,

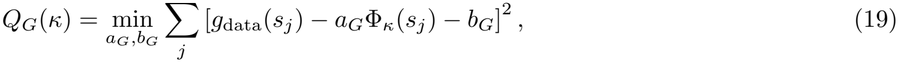

and 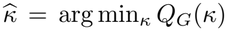. At the behavioral stage, 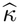 is held fixed and only the gain and offset of the behavioral readout are estimated

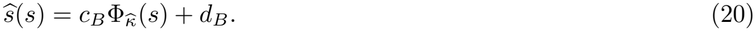

This is the affine behavioral readout introduced above, expressed in the normalized coordinate: if 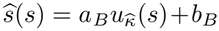, then 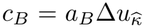 and 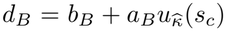. The neural fit fixes 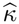; behavior supplies only an independent gain and offset. Agreement with 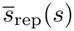 therefore tests the nonlinear shape inferred from neural geometry.

The normalized solution has two asymptotic limits separated by a continuous crossover. At small *κ*, boundary distortions can be unresolved at the sampling resolution and measurement precision, yielding an approximately affine shape. Intermediate *κ* gives resolved nonlinear shapes, whereas large *κ* gives a limiting sigmoid with overlapping boundary effects. Accordingly, *Q*_*G*_(*κ*) approaches plateaus in both limits; the data may favor either plateau or a finite interior minimum. For mathematical details see the Supplemental Material. The 20 networks were classified as small-*κ* (4), intermediate-*κ* (9), and large-*κ* (7) (Fig. 3 (g–i), Supplemental Fig. S3 and Supplemental Table SI). Plateaus identify limiting shapes rather than unique finite parameters.

Representative neural fits and conditional behavioral predictions are shown in Fig. 3 (a–c) and (d–f), respectively (full lines). The geometry-selected shapes capture the systematic dependence of mean reproduction on stimulus interval, with independent neural and behavioral gains and offsets. All behavioral predictions, neural fits, and loss profiles are shown in Supplemental Figs. S1, S2, and S3, respectively.

### Relation to Bayesian estimation

Bayesian observer models explain central-tendency biases by combining noisy measurements with prior knowledge [5, 7, 18]. For comparison with the exact variational solution, we consider two

Bayesian priors over the trained interval range: a continuous bounded-uniform prior [7], corresponding to the coarse-grained weighting used in the variational model,

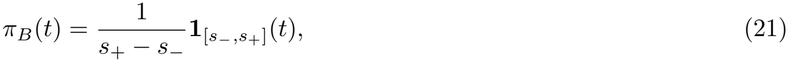

and a discrete-uniform prior over the *J* = 11 trained intervals, corresponding to the actual training ensemble. The following definitions apply separately to each prior; the prior index is suppressed for brevity. Measurements were modeled by a Gaussian likelihood *p*(*m* | *t*; *w*_*m*_) with scalar variance 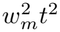. The Bayesian least-squares estimator *f*_BLS_, for a measurement *m*, is the mean of the corresponding posterior distribution and its mean response at physical stimulus *s* is

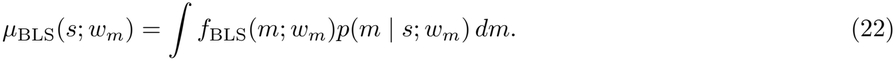

The integrals defining the BLS mean-response curve were evaluated numerically. The affine-invariant Bayesian shape was defined as

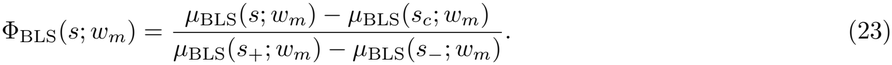

For each network, we estimate *w*_*m*_ from the neural geometric data by profiling out the neural gain and offset,

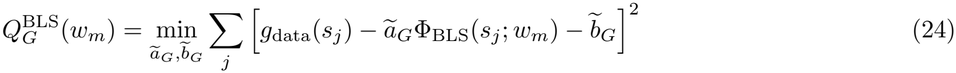

where *ã*_*G*_ and 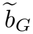 are affine parameters. Then we compute 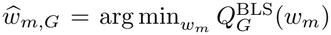. Holding this parameter fixed, the BLS prediction of the mean behavioral response is

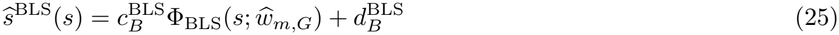

where 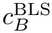 and 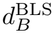 are the behavioral gain and offset.

Figure 4 compares in-sample geometric root-mean-square errors (RMSEs) across all 20 networks. The BLS discrete (BLS-D) and continuous (BLS-C) models have lower errors than the variational model in 9 and 11 networks, respectively. Supplemental Figs. S4–S6 show the BLS behavioral predictions, neural fits, and loss profiles, respectively; Supplemental Table SI lists the geometry-selected parameters and RMSEs of all three models.

**FIG. 4.**
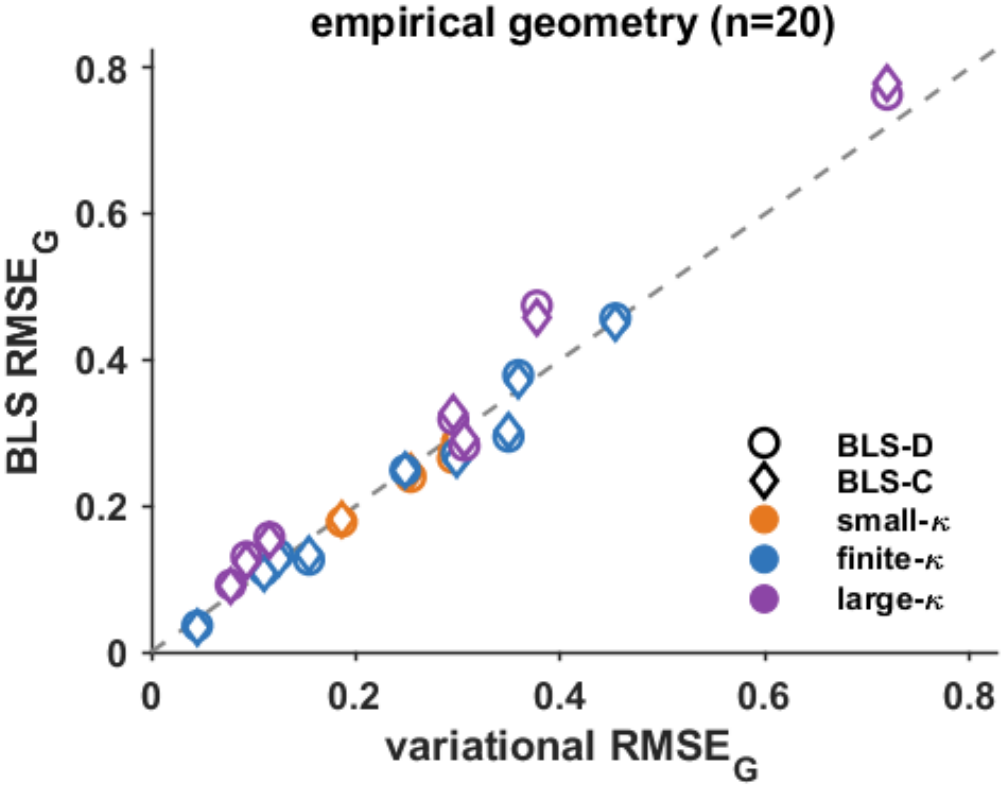
Variational and BLS fits to neural geometry. In-sample geometric RMSEs for 20 networks: variational model (horizontal) and BLS (vertical). Circles denote the discrete-uniform prior (BLS-D); diamonds, the continuous-uniform prior (BLS-C). Colors indicate small-*κ* (orange), intermediate-*κ* (blue), and large-*κ* (purple) variational regimes. Each model’s shape parameter, gain, and offset are fitted independently to the same neural data. The gray dashed identity line indicates equal geometric RMSEs.

Separately, we compared the model shapes by minimizing the affine-invariant discrepancy defined as

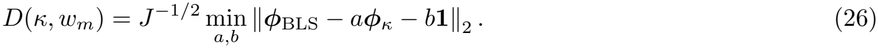

Here ***ϕ***_*κ*_ and ***ϕ***_BLS_ are *J*-dimensional column vectors with components given in Eqs. (16) and (23), respectively, and **1** is a vector of ones. Because Φ_BLS_ is normalized by the Bayesian response range, *D* is expressed as a fraction of that range. For each geometry-selected *κ*, this comparison uses only theoretical curves and defines a shape-matching parameter 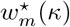, distinct from 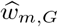 . The minimized discrepancies for the two BLS models are denoted as *D* and *D*_*D*_. The two families were very close. For intermediate-*κ* and large-*κ* networks, the median 100*D*_*C*_ is below 1% and approximately 2%, respectively; the corresponding medians of 100*D*_*D*_ are 1.26% and 2.29%. Shape-matching parameters and continuous-prior discrepancies are listed in Supplemental Table SII.

This correspondence is boundary-driven. In the variational model, the boundary conditions propagate through the Euler–Lagrange equation; in the BLS model, truncation of the posterior by the bounded prior generates a similar sigmoidal mean response [7]. Thus, distinct mathematical operations produce closely related affine-invariant geometries.

## Discussion

The proposed variational principle relates learned neural geometry to stimulus weighting, task fidelity, and smoothness. In interval-reproduction networks, its boundary-layer regimes organize the observed geometries, and the shape inferred from neural activity captures the nonlinear component of mean behavior through an affine readout. The framework may also extend to delayed-comparison tasks, where distortions of a remembered stimulus can bias subsequent choices. Such an extension would connect the present account to contraction biases and memory representations studied in experimental and modeling work [11, 12, 18–21] .

Our results suggest a principle through which Bayesian-like behavior may arise from the organization of learned representations. BLS specifies an optimal estimator for a given prior, likelihood, and loss, whereas the variational principle determines a latent representation. Their closely matching affine-normalized shapes suggest a geometric route to Bayesian-like behavior.

## Acknowledgments

The authors used ChatGPT (OpenAI) to improve the language and assess consistency throughout the text. The authors subsequently revised all AI-assisted content and assume full responsibility for the final manuscript.

