## Supplementary Information for "A Variational Principle for the Emergence of Neural Geometry Underlying Bayesian-Like Behavior"

*Departamento de Física Teórica, Universidad Autónoma de Madrid,*
*28049 Madrid, Spain*

### CONTENTS

**Supplemental Text** **1**

**1 Exact bounded-uniform solutions** **1**

**2 Affine-invariant shapes and three boundary-layer regimes** **2**

**3 Bayesian models and numerical evaluation** **4**

**Supplemental Figures** **6**

Fig. S1 Conditional variational predictions for all networks . . . . . . 6
Fig. S2 Neural geometry and variational fits for all networks . . . . . . 7
Fig. S3 Profiled variational geometric losses . . . . . . 8
Fig. S4 Conditional BLS predictions for all networks . . . . . . 9
Fig. S5 Neural geometry and BLS fits for all networks . . . . . . 10
Fig. S6 Profiled BLS geometric losses . . . . . . 11

**Supplemental Tables** **12**

Table SI Variational and BLS fits to neural geometry . . . . . . 12
Table SII Theoretical shape correspondence with BLS . . . . . . 13

**References** **13**

### Supplemental Text

### 1 Exact bounded-uniform solutions

*Constant inverse-fidelity scale.* For a bounded-uniform stimulus-weighting density on $[s_-, s_+]$, the score vanishes in the interior of the support. For constant inverse fidelity, $\Sigma_G(s) = \Sigma_0$, the Euler–Lagrange equation and its natural boundary conditions are

$$u(s) - s - \ell^2 u''(s) = 0, \qquad u'(s_-) = u'(s_+) = 0, \tag{S1}$$

where $\ell^2 = \lambda\Sigma_0$. Define the midpoint and width of the support as

$$\bar{s} = \frac{s_- + s_+}{2}, \qquad \Delta s = s_+ - s_-. \tag{S2}$$

The exact solution satisfying the boundary conditions is

$$u(s) = s - \ell\, \frac{\sinh[(s - \bar{s})/\ell]}{\cosh[\Delta s/(2\ell)]}. \tag{S3}$$

*; Also at Centro de Investigación Avanzada en Física Fundamental, Universidad Autónoma de Madrid, 28049 Madrid, Spain.

When $\Delta s/(2\ell) \gg 1$, Eq. (S3) reduces to

$$u(s) \simeq s + \ell \exp\left[-\frac{s - s_-}{\ell}\right] - \ell \exp\left[-\frac{s_+ - s}{\ell}\right]. \tag{S4}$$

Consequently, $u(s) \simeq s$ at points whose distance from either endpoint is large compared with $\ell$, in agreement with the weak-distortion prediction [Eq. (8) of the main text]. However, every finite-$\ell$ solution has zero slope at the two endpoints. The regular weak-distortion expansion therefore misses the boundary layers required by the natural boundary conditions.

*Scalar inverse-fidelity scale.* For time intervals $s > 0$, we take

$$\Sigma_G(s) = w_G^2 s^2, \qquad \kappa^2 = \lambda w_G^2. \tag{S5}$$

The exact variational coordinate obeys

$$u_\kappa(s) - s - \kappa^2 s^2 u_\kappa''(s) = 0, \qquad u_\kappa'(s_-) = u_\kappa'(s_+) = 0. \tag{S6}$$

The general solution is

$$u_\kappa(s) = s + A_\kappa s^{r_+} + B_\kappa s^{r_-}, \qquad r_\pm = \frac{1}{2}\left(1 \pm \sqrt{1 + \frac{4}{\kappa^2}}\right), \tag{S7}$$

where $A_\kappa$ and $B_\kappa$ are determined by the two Neumann boundary conditions.

For numerical evaluation, we used the form

$$u_\kappa(s) = s + C_-\left(\frac{s}{s_-}\right)^{r_-} + C_+\left(\frac{s}{s_+}\right)^{r_+}. \tag{S8}$$

The coefficients $C_-$ and $C_+$ satisfy

$$\frac{r_-}{s_-} C_- + \frac{r_+}{s_+}\left(\frac{s_-}{s_+}\right)^{r_+ - 1} C_+ = -1, \tag{S9}$$

$$\frac{r_-}{s_-}\left(\frac{s_+}{s_-}\right)^{r_- - 1} C_- + \frac{r_+}{s_+} C_+ = -1. \tag{S10}$$

#### 2 Affine-invariant shapes and three boundary-layer regimes

Overall contraction and nonlinear geometric distortion must be distinguished. An affine stimulus–response relation with slope smaller than one can compress reproduced intervals, whereas nonlinear distortion involves a stimulus-dependent slope. Because the neural observation model and behavioral readout have independent gains and offsets, overall contraction alone does not identify $\kappa$. The neural fit constrains $\kappa$ through the nonlinear shape that remains after affine alignment.

Consider the bounded-uniform scalar model with $\Sigma_G(s) = w_G^2 s^2$ and $\kappa^2 = \lambda w_G^2$. As in the main text, we define

$$\begin{aligned} \Phi_\kappa(s) &= \frac{u_\kappa(s) - u_\kappa(s_c)}{\Delta u_\kappa}, \\ \Delta u_\kappa &= u_\kappa(s_+) - u_\kappa(s_-), \end{aligned} \tag{S11}$$

Thus $\Phi_\kappa(s_c) = 0$ and $\Phi_\kappa(s_+) - \Phi_\kappa(s_-) = 1$. The normalization removes any nonzero affine gain and offset.

The three-regime description follows from two distinct asymptotic limits and the crossover between them. As $\kappa \to 0$, the endpoint propagation scales $\ell_\pm = \kappa s_\pm$ vanish and the normalized solution approaches an affine function. At intermediate values, changing $\kappa$ changes the extent and overlap of boundary distortions. As $\kappa \to \infty$, boundary effects span the full interval and the normalized shape approaches a fixed nonlinear limit. These are regimes of one continuous solution family, not separate solution branches; their empirical separation depends on stimulus sampling and measurement precision.

To connect these shapes with parameter estimation, we use the profiled geometric loss defined in the main text,

$$Q_G(\kappa) = \min_{a_G, b_G} \sum_j \left[g_{\text{data}}(s_j) - a_G \Phi_\kappa(s_j) - b_G\right]^2. \tag{S12}$$

Its two asymptotic plateaus follow from convergence of the normalized shapes. Whether the data favor either plateau or a finite interior minimum is an empirical question.

*Small-$\kappa$ regime: unresolved boundary layers.—* Away from the endpoints, the solution approaches $u_\kappa(s) = s$. For narrow, well-separated boundary layers, the leading composite approximation is

$$\begin{aligned} u_\kappa(s) \simeq s + \kappa s_- \exp\left[-\frac{s - s_-}{\kappa s_-}\right] \\ - \kappa s_+ \exp\left[-\frac{s_+ - s}{\kappa s_+}\right]. \end{aligned} \tag{S13}$$

Consequently, with $\Delta s = s_+ - s_-$,

$$\Phi_\kappa(s) \longrightarrow \Phi_0(s) = \frac{s - s_c}{\Delta s}, \qquad Q_G(\kappa) \longrightarrow Q_{G,0}. \tag{S14}$$

Here $Q_{G,0}$ is the least-squares loss of an affine fit to the neural data. For every $\kappa > 0$, however, $\Phi'_\kappa(s_-) = \Phi'_\kappa(s_+) = 0$: the convergence is singular in the endpoint derivatives. When $\ell_\pm$ are much smaller than the spacing between sampled stimuli, the endpoint transition is not resolved. Indistinguishability from an affine fit additionally requires the residual shape deviations, including sampled endpoint displacements, to fall below measurement precision. A preferred small-$\kappa$ plateau therefore identifies an approximately affine shape, rather than a particular nonzero boundary-layer width.

*Intermediate-$\kappa$ regime: resolved changes of shape.—* Here the sampled geometry resolves changes in $\Phi_\kappa$ as $\kappa$ varies. The exact solution is increasing, with zero endpoint slopes, and satisfies

$$\Phi''_\kappa(s) = \frac{u_\kappa(s) - s}{\kappa^2 s^2 \Delta u_\kappa}. \tag{S15}$$

It is convex below its unique crossing of $u_\kappa(s) = s$ and concave above it, producing a sigmoid with endpoint flattening. For separated layers, $\ell_+ > \ell_-$ implies a broader upper boundary region; increasing $\kappa$ extends their influence into the interior. If the data distinguish a finite shape from both limiting shapes, $Q_G$ can have a localized interior minimum at $\widehat{\kappa}$. Such a minimum supports estimation of a finite propagation scale, provided that its depth and width are resolved relative to measurement uncertainty. A finite minimum is not guaranteed merely by taking $\kappa$ between its asymptotic limits.

*Large-$\kappa$ regime: a limiting global sigmoid.—* The unnormalized solution has the expansion

$$u_\kappa(s) = U_0 + \kappa^{-2} v(s) + O(\kappa^{-4}). \tag{S16}$$

Substitution into the scalar boundary-value problem gives

$$s^2 v''(s) = U_0 - s, \qquad v'(s_-) = v'(s_+) = 0. \tag{S17}$$

Integrating $v''$ across the interval and imposing both endpoint conditions determines

$$U_0 = \frac{s_- s_+ \log(s_+/s_-)}{\Delta s}, \qquad s_- < U_0 < s_+. \tag{S18}$$

The derivative is

$$v'(s) = U_0 \left(\frac{1}{s_-} - \frac{1}{s}\right) - \log\left(\frac{s}{s_-}\right). \tag{S19}$$

The additive constant in $v$ is fixed by $\int_{s_-}^{s_+} v(s) s^{-2}\, ds = 0$ for the expansion above, but cancels from the normalized shape. Since $v''(s)$ changes sign at $U_0$, $v'(s)$ increases from zero, reaches its maximum there, and decreases to zero at $s_+$. Thus $v$ defines an increasing, generally asymmetric sigmoid.

Although the raw dynamic range vanishes, normalization retains this shape:

$$\begin{aligned}\Delta u_\kappa &= \kappa^{-2}\big[v(s_+) - v(s_-)\big] + O(\kappa^{-4}),\\ \Phi_\kappa(s) &= \Phi_\infty(s) + O(\kappa^{-2}),\\ \Phi_\infty(s) &= \frac{v(s) - v(s_c)}{v(s_+) - v(s_-)}.\end{aligned} \tag{S20}$$

It follows that $Q_G(\kappa) \to Q_{G,\infty}$, the profiled loss for the template $\Phi_\infty$. The normalized gains $a_G$ and $c_B$ need not diverge: the shrinking range of $u_\kappa$ has already been removed. A preferred large-$\kappa$ plateau identifies this limiting shape, rather than a finite value of $\kappa$ or a collapse of the measured neural representation.

*Consequences for behavior.*— With $\widehat{\kappa}$ selected from neural geometry, the conditional behavioral prediction remains

$$\widehat{s}(s) = c_B\Phi_{\widehat{\kappa}}(s) + d_B. \tag{S21}$$

In the small-$\kappa$ limit this is affine, with slope $c_B/\Delta s$, so overall contraction can persist without resolved nonlinear distortion. The intermediate and large-$\kappa$ regimes instead predict the corresponding nonlinear shapes under an affine readout. For either preferred plateau, a numerical search boundary should be reported as an asymptotic classification, or as a one-sided constraint when supported by an uncertainty analysis, rather than as a uniquely identified finite estimate.

### 3 Bayesian models and numerical evaluation

*Continuous bounded-uniform prior.* This Bayesian model has been considered in [1]; it is included here for the sake of completeness. For the continuous bounded-uniform prior,

$$\pi_B(t) = \frac{1}{s_+ - s_-}\mathbf{1}_{[s_-,s_+]}(t), \tag{S22}$$

the Bayesian least-squares estimator is

$$f_{\mathrm{BLS}}(m; w_m) = \frac{\displaystyle\int_{s_-}^{s_+} t\, p(m \mid t; w_m)\, dt}{\displaystyle\int_{s_-}^{s_+} p(m \mid t; w_m)\, dt}. \tag{S23}$$

Although $\pi_B(t)$ has the same functional form as the stimulus weighting [Eq. (12) of the main text], it plays a different role: $\pi_B$ is the prior of the Bayesian observer, whereas $p(s)$ weights the variational learning objective. The analogous construction for the literal discrete-uniform prior over the $J = 11$ trained intervals is defined below.

The corresponding mean-response curve, conditional on the physical stimulus $s$, is

$$\mu_{\mathrm{BLS}}(s; w_m) = \int_{-\infty}^{\infty} f_{\mathrm{BLS}}(m; w_m)\, p(m \mid s; w_m)\, dm. \tag{S24}$$

The scalar Gaussian measurement likelihood is

$$p(m \mid t; w_m) = \frac{1}{\sqrt{2\pi}\, w_m t}\exp\left[-\frac{(m-t)^2}{2w_m^2 t^2}\right]. \tag{S25}$$

The integrals over the prior support in Eq. (S23) were evaluated using Gauss–Legendre quadrature. The average over the measurement distribution in Eq. (S24) was evaluated using Gauss–Hermite quadrature. Doubling the quadrature orders did not change the reported values at the displayed precision.

The affine-invariant Bayesian shape is defined as

$$\Phi_{\mathrm{BLS}}(s; w_m) = \frac{\mu_{\mathrm{BLS}}(s; w_m) - \mu_{\mathrm{BLS}}(s_c; w_m)}{\mu_{\mathrm{BLS}}(s_+; w_m) - \mu_{\mathrm{BLS}}(s_-; w_m)}. \tag{S26}$$

*Discrete-uniform training prior.* Because the networks were trained on $J = 11$ discrete stimulus values, we also considered the discrete-uniform prior

$$\pi_D(t) = \frac{1}{J}\sum_{j=1}^{J}\delta(t - s_j). \tag{S27}$$

For this prior, the Bayesian least-squares estimator is

$$f_D(m; w_m) = \frac{\sum_{j=1}^{J} s_j \, p(m \mid s_j; w_m)}{\sum_{j=1}^{J} p(m \mid s_j; w_m)}. \tag{S28}$$

Its conditional mean-response curve is

$$\mu_D(s; w_m) = \int_{-\infty}^{\infty} f_D(m; w_m) \, p(m \mid s; w_m) \, dm. \tag{S29}$$

### Supplemental Figures

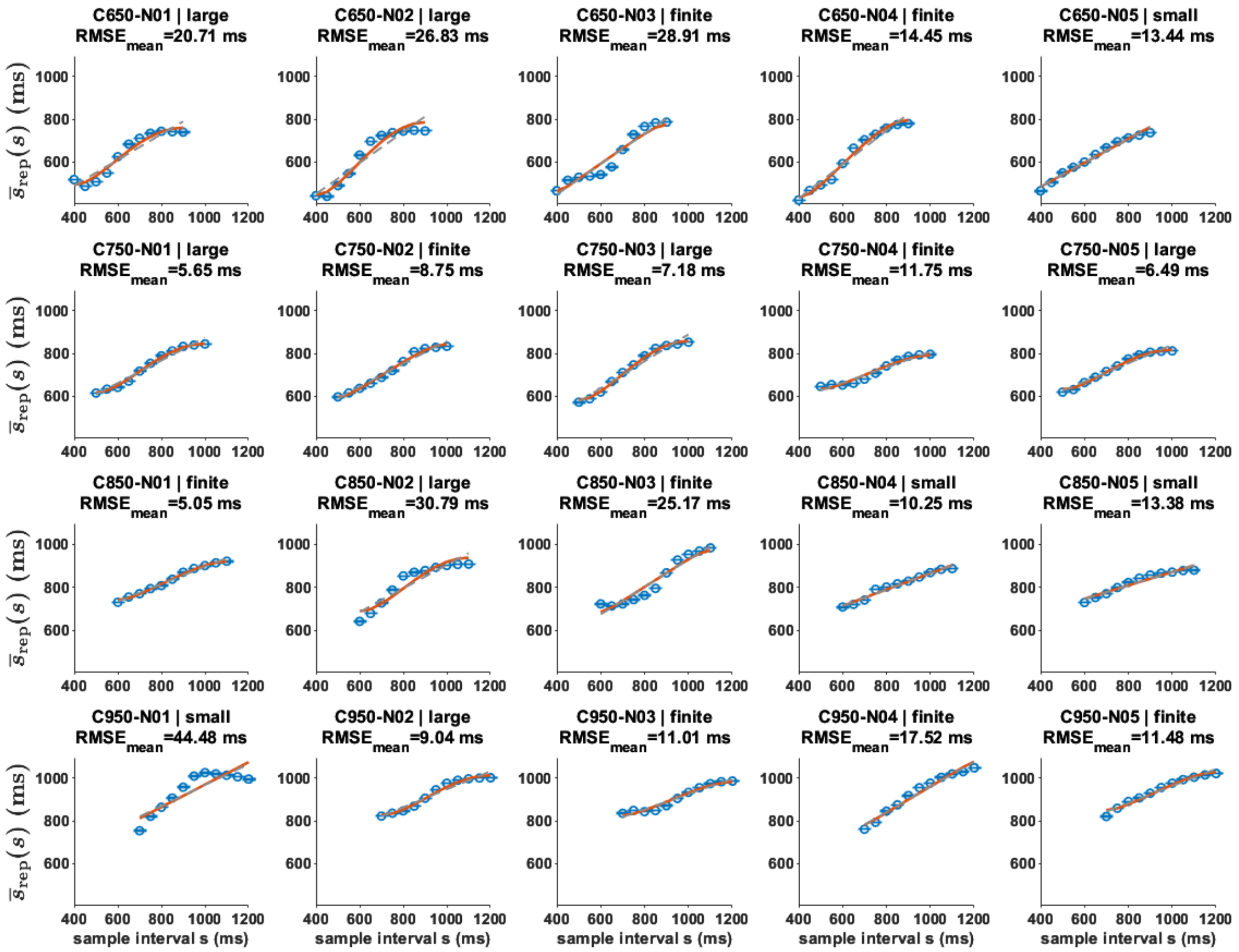


Figure S1: **Time-interval reproduction and conditional variational predictions for all 20 networks.** Each panel shows one network. Rows correspond to training ensembles centered at $C = 650$, 750, 850, and 950 ms, from top to bottom, with five networks per center. Open circles show the mean reproduced interval $\bar{s}_{\mathrm{rep}}(s)$ at each sample interval $s$. Orange curves show $\widehat{s}(s) = c_B \Phi_{\widehat{\kappa}}(s) + d_B$, where $\widehat{\kappa}$ is selected from the neural geometric data and held fixed when fitting the behavioral gain $c_B$ and offset $d_B$. Gray dashed lines show the affine null model fitted to the reproduction means. Panel titles identify each network and report the root-mean-square error between the observed stimulus-wise mean reproduced intervals and the conditional model predictions, in milliseconds ($\mathrm{RMSE}_{\mathrm{mean}}$).

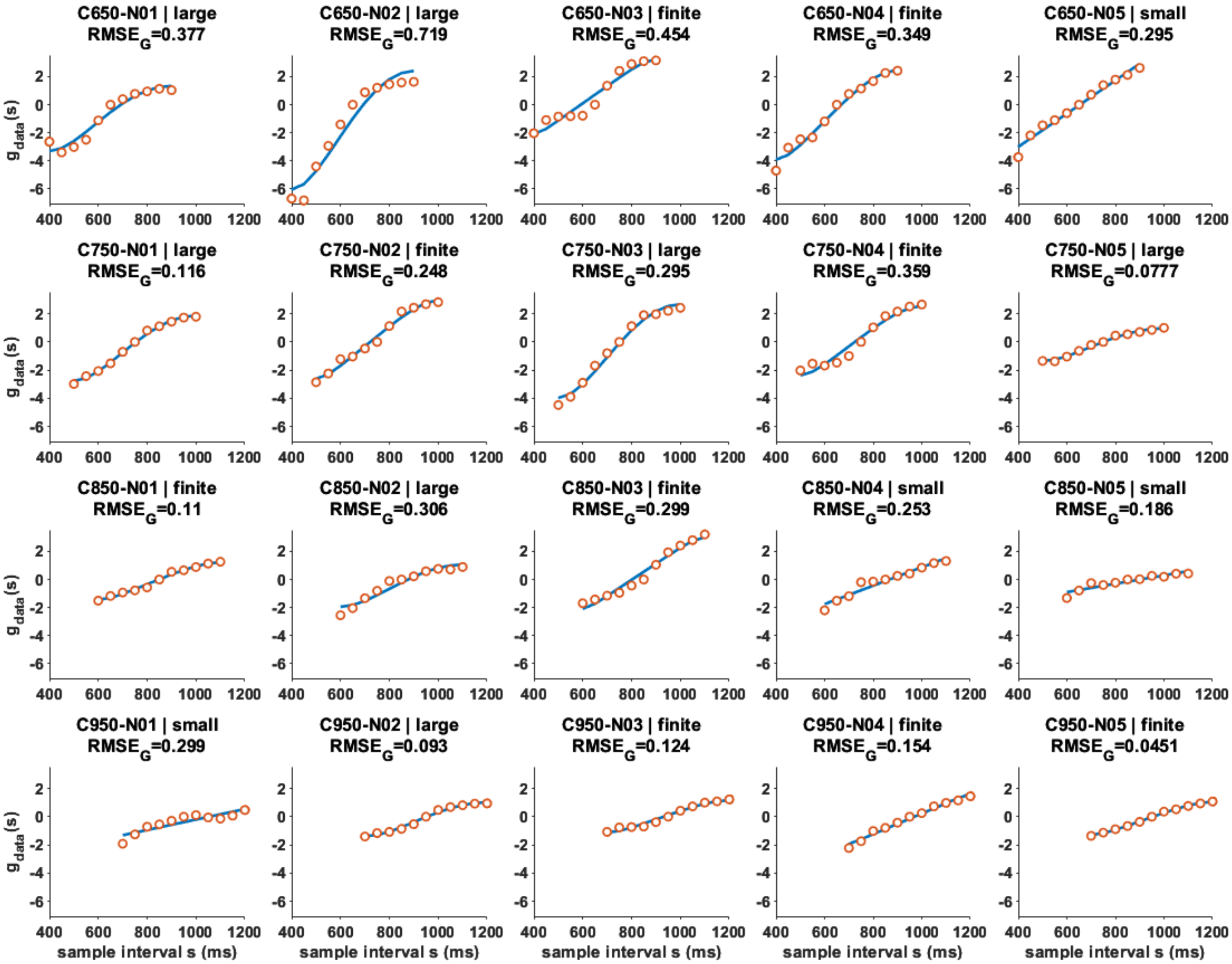


Figure S2: **Empirical neural geometry and variational fits for all 20 networks.** Panels follow the same network ordering as Fig. S1, with rows corresponding to training ensemble centers $C$ = 650, 750, 850, and 950 ms. Open circles show the empirical geometric coordinate $g_{\text{data}}(s)$ obtained from signed distances to a reference trajectory. Blue solid curves show the fitted observation model $a_G \Phi_{\widehat{\kappa}}(s) + b_G$. For each network, the affine parameters $a_G$ and $b_G$ are optimized at every candidate $\kappa$, and $\widehat{\kappa}$ is selected by minimizing the resulting profiled geometric loss $Q_G(\kappa)$. Panel titles identify the network and report the geometric root-mean-square error across sampled intervals, in the units of $g_{\text{data}}$. These neural fits determine the latent shapes used for the conditional behavioral predictions in Fig. S1.

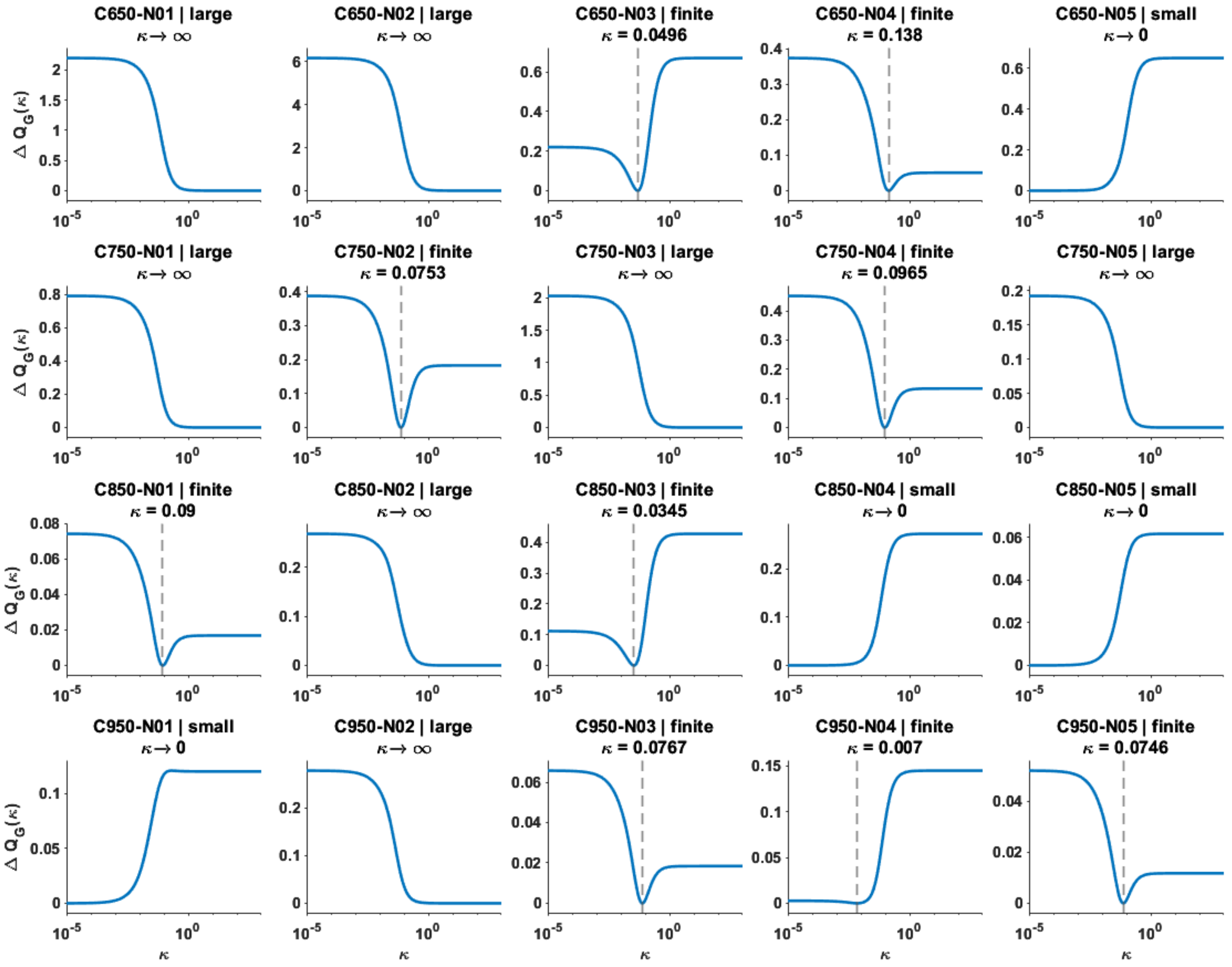


Figure S3: **Profiled geometric losses across networks.** Blue curves show $\Delta Q_G(\kappa) = Q_G(\kappa) - \min_{\kappa'} Q_G(\kappa')$ against $\kappa$ on a logarithmic scale, with neural gain and offset optimized at each $\kappa$. Panels follow the ordering of Supplemental Fig. S1. Titles identify small-$\kappa$ plateaus (4 networks), finite interior minima (9), and large-$\kappa$ plateaus (7). For finite minima, the numerical values of $\kappa$ displayed in the panel titles correspond to the geometry-selected $\widehat{\kappa}$, marked by gray dashed vertical lines. For plateau regimes, the labels $\kappa \to 0$ and $\kappa \to \infty$ denote asymptotic limits rather than identifiable finite parameter estimates.

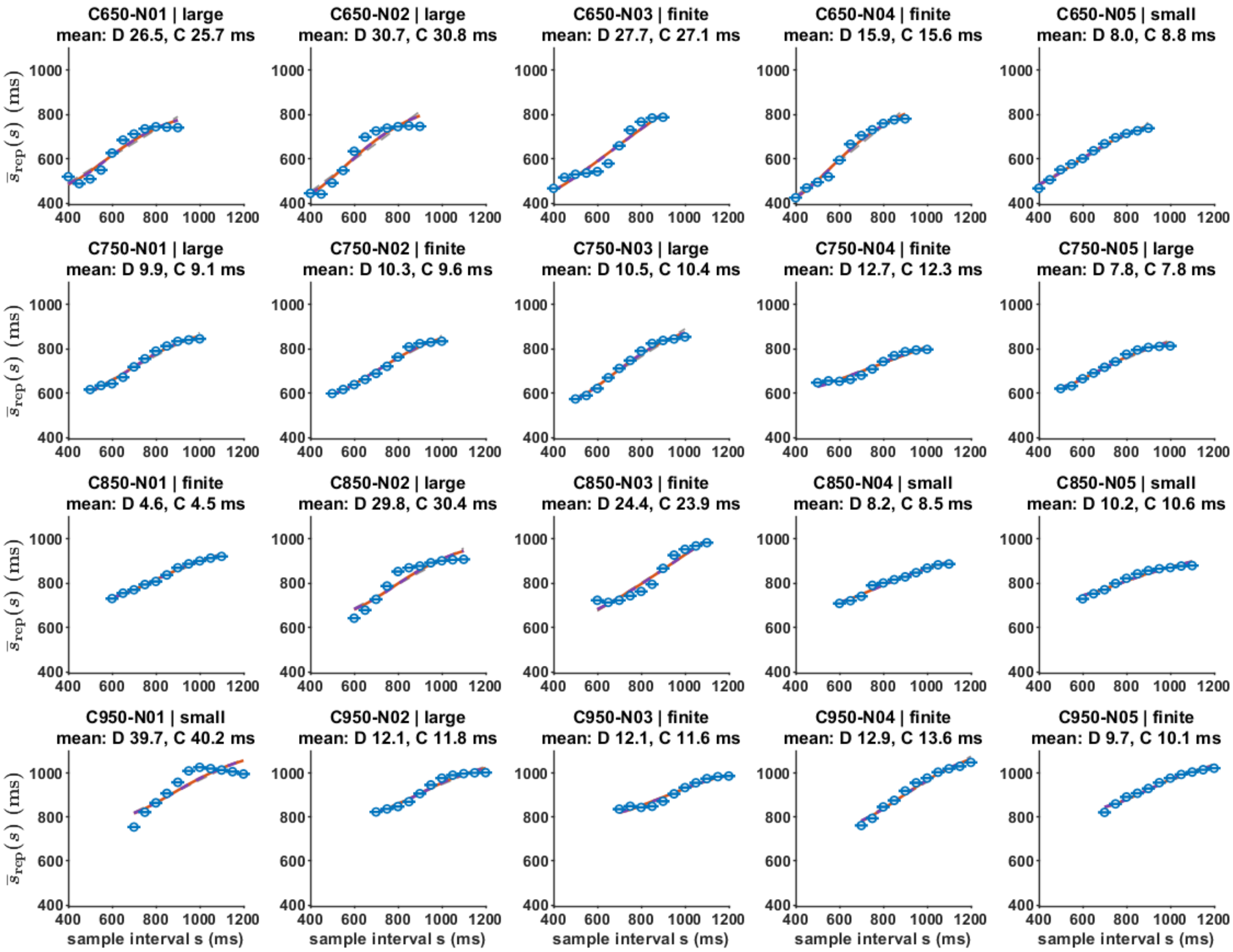


Figure S4: **Time-interval reproduction and conditional Bayesian least-squares (BLS) predictions for all 20 networks.** Panels follow the same network ordering as Fig. S1, with rows corresponding to training ensemble centers $C = 650$, 750, 850, and 950 ms. Open blue circles show the observed mean reproduced intervals $\overline{s}_{\mathrm{rep}}(s)$ at each sample interval $s$, with error bars indicating the standard error of the mean when available. Orange solid curves and purple dashed curves show the conditional predictions obtained with BLS using discrete-uniform (BLS-D) and continuous-uniform (BLS-C) priors, respectively. For each model, the parameter $\widehat{w}_m$ is selected from neural geometry and held fixed when fitting the affine behavioral readout. Only the behavioral gain and offset are fitted to the stimulus-wise reproduction means. Gray dashed curves indicate the affine null model fitted to the same means. Panel titles identify the network and its variational regime and report the root-mean-square errors of the stimulus-wise means for BLS-D and BLS-C, in milliseconds. These errors quantify agreement with the displayed means rather than trial-level reproduction errors.

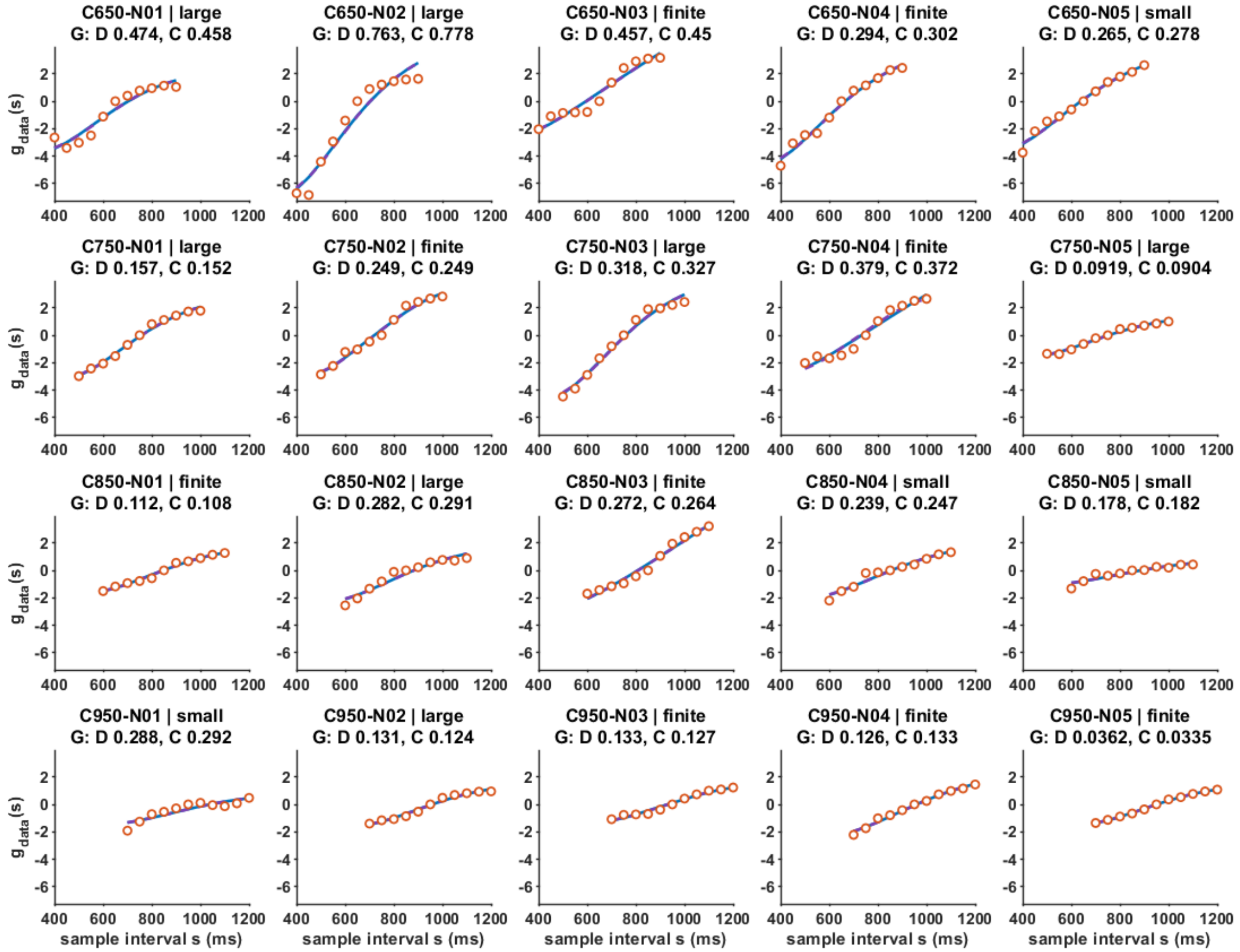


Figure S5: **Empirical neural geometry and Bayesian least-squares (BLS) fits for all 20 networks.** Panels follow the same network ordering as Fig. S1, with rows corresponding to training ensembles centered at $C = 650$, 750, 850, and 950 ms and five networks per center. Open orange circles show the empirical neural geometric coordinates $g_{\text{data}}(s)$. Blue solid curves and purple dashed curves show the fitted geometric coordinates obtained with BLS using discrete-uniform (BLS-D) and continuous-uniform (BLS-C) priors, respectively. For each network, the BLS parameter $w_m$ and the affine geometric gain and offset are selected by fitting the neural geometric data. Panel titles identify the network and its variational regime and report the geometric root-mean-square errors $\text{RMSE}_{G,D}$ and $\text{RMSE}_{G,C}$ for the two BLS models. All panels use common axis limits to facilitate comparison across networks.

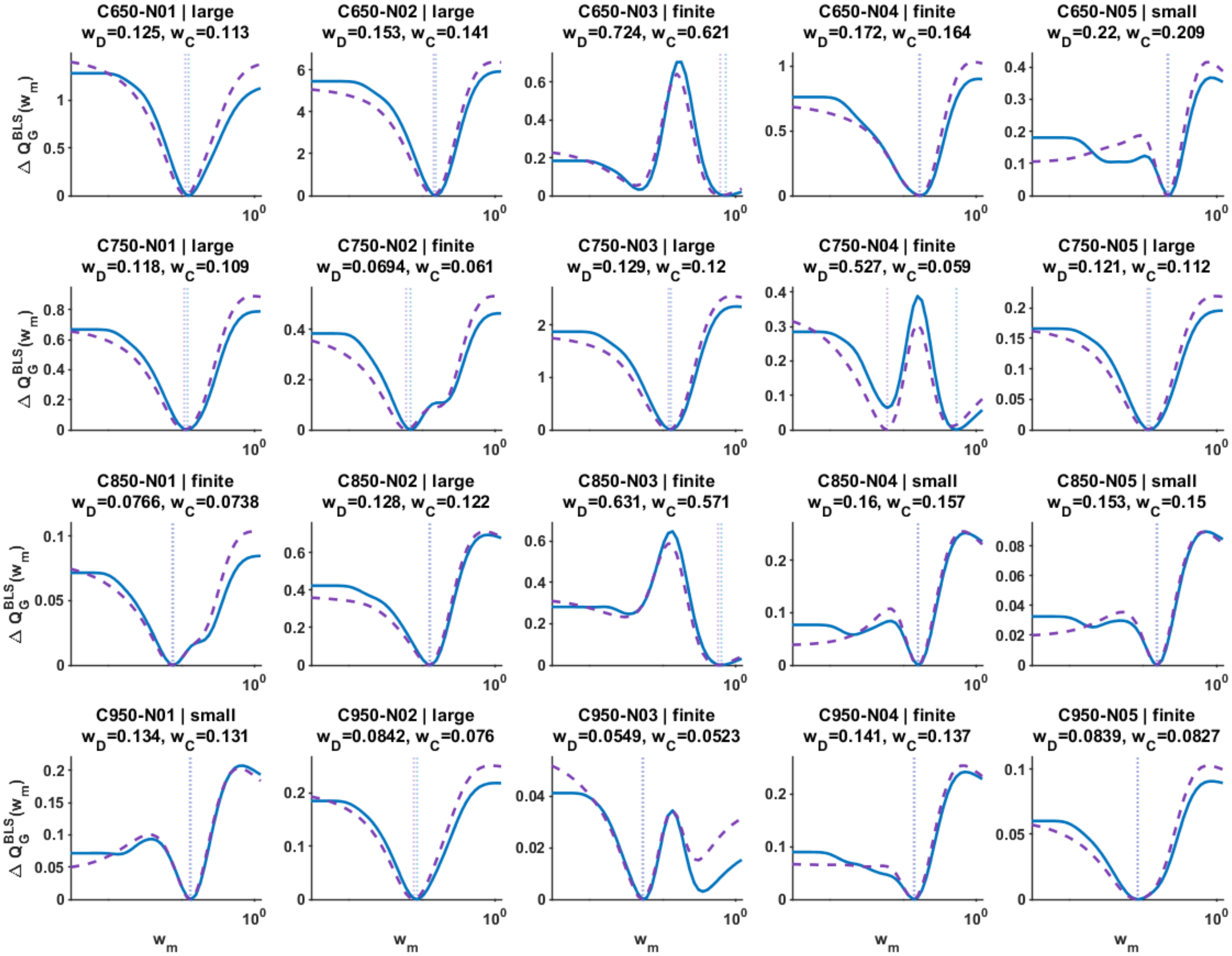


Figure S6: **Profiled geometric losses for Bayesian least-squares (BLS) fits across all 20 networks.** Each panel shows the excess BLS geometric loss $\Delta Q_G^{\mathrm{BLS}}(w_m)$ as a function of parameter $w_m$ on a logarithmic horizontal axis, with separate profiles for the discrete- and continuous-uniform priors. Blue solid curves correspond to the discrete-uniform prior (BLS-D), and purple dashed curves correspond to the continuous-uniform prior (BLS-C). For each prior, the loss is computed as the sum of squared residuals between the empirical neural geometric coordinates $g_{\mathrm{data}}(s)$ and the fitted BLS coordinates, with the affine geometric gain and offset optimized at each $w_m$. Each profile is shifted by its own minimum over the evaluated parameter grid. Vertical dotted lines indicate the corresponding geometry-selected parameters $\widehat{w}_{m,D}$ and $\widehat{w}_{m,C}$ obtained from the original BLS fits. Panels follow the same network ordering as Fig. S1. The vertical limits are adjusted independently for each panel to make differences in profile shape visible. The profiles are evaluated numerically on a finite logarithmic grid; therefore, their minima should be interpreted at the resolution of that grid.

### Supplemental Tables

Table SI: **Network-by-network variational and BLS fits to neural geometry.** Networks are grouped according to the three regimes of the variational model: small-$\kappa$ plateau, finite-$\kappa$ minimum, and large-$\kappa$ plateau. The parameter $\widehat{\kappa}$ is selected from neural geometry, with $\kappa \to 0$ and $\kappa \to \infty$ indicating asymptotic plateaus rather than identifiable finite values. The quantities $\widehat{w}_{m,D}$ and $\widehat{w}_{m,C}$ are the BLS parameters selected from neural geometry using discrete- and continuous-uniform priors, respectively. The last three columns report the in-sample geometric root-mean-square errors ($RMSE_G$) for the variational model ($V$), discrete BLS ($D$), and continuous BLS ($C$). All errors are evaluated against the same empirical neural geometric coordinates $g_{\text{data}}(s)$, using identical stimulus sets and units. Network identifiers include the training-ensemble center (in ms).

| Network | $\widehat{\kappa}$ | $\widehat{w}_{m,D}$ | $\widehat{w}_{m,C}$ | $\text{RMSE}_{G,V}$ | $\text{RMSE}_{G,D}$ | $\text{RMSE}_{G,C}$ |
|---|---|---|---|---|---|---|
| Small-$\kappa$ plateau (4 networks) | | | | | | |
| C650-N05 | $\kappa \to 0$ | 0.220 | 0.209 | 0.295 | 0.265 | 0.278 |
| C850-N04 | $\kappa \to 0$ | 0.160 | 0.157 | 0.254 | 0.239 | 0.247 |
| C850-N05 | $\kappa \to 0$ | 0.153 | 0.150 | 0.186 | 0.178 | 0.182 |
| C950-N01 | $\kappa \to 0$ | 0.134 | 0.131 | 0.299 | 0.288 | 0.292 |
| Finite-$\kappa$ minimum (9 networks) | | | | | | |
| C650-N03 | 0.0496 | 0.724 | 0.621 | 0.454 | 0.457 | 0.451 |
| C650-N04 | 0.138 | 0.172 | 0.164 | 0.349 | 0.294 | 0.302 |
| C750-N02 | 0.0753 | 0.0694 | 0.0610 | 0.248 | 0.249 | 0.249 |
| C750-N04 | 0.0965 | 0.527 | 0.0590 | 0.359 | 0.379 | 0.372 |
| C850-N01 | 0.0900 | 0.0766 | 0.0738 | 0.110 | 0.112 | 0.108 |
| C850-N03 | 0.0345 | 0.631 | 0.571 | 0.299 | 0.272 | 0.264 |
| C950-N03 | 0.0767 | 0.0549 | 0.0523 | 0.124 | 0.133 | 0.127 |
| C950-N04 | 0.00700 | 0.141 | 0.137 | 0.154 | 0.126 | 0.133 |
| C950-N05 | 0.0746 | 0.0839 | 0.0827 | 0.0451 | 0.0362 | 0.0335 |
| Large-$\kappa$ plateau (7 networks) | | | | | | |
| C650-N01 | $\kappa \to \infty$ | 0.125 | 0.113 | 0.377 | 0.475 | 0.458 |
| C650-N02 | $\kappa \to \infty$ | 0.153 | 0.141 | 0.719 | 0.763 | 0.779 |
| C750-N01 | $\kappa \to \infty$ | 0.118 | 0.109 | 0.116 | 0.157 | 0.152 |
| C750-N03 | $\kappa \to \infty$ | 0.129 | 0.120 | 0.295 | 0.318 | 0.327 |
| C750-N05 | $\kappa \to \infty$ | 0.121 | 0.112 | 0.0777 | 0.0919 | 0.0904 |
| C850-N02 | $\kappa \to \infty$ | 0.128 | 0.122 | 0.306 | 0.282 | 0.291 |
| C950-N02 | $\kappa \to \infty$ | 0.0842 | 0.0760 | 0.0930 | 0.131 | 0.124 |

Table SII: **Theoretical shape correspondence between the variational and BLS models.** Networks are grouped according to the three variational regimes: small-$\kappa$ plateau, finite-$\kappa$ minimum, and large-$\kappa$ plateau. The quantities $w^\star_{m,D}$ and $w^\star_{m,C}$ are the theoretical shape-matching parameters for BLS with discrete- and continuous-uniform priors, respectively. The final column reports the continuous-prior shape discrepancy $100D_C$, expressed as a percentage, after removing affine scale and offset differences. For the small-$\kappa$ networks, the variational shapes are approximately affine, making the shape-matching parameters poorly constrained and the comparison of nonlinear distortions less informative. Consequently, $100D_C$ is not reported for these networks (dashes). Network identifiers include the training-ensemble center C (in ms).

| Network | $w^\star_{m,D}$ | $w^\star_{m,C}$ | $100D_C$ |
|---|---|---|---|
| | Small-$\kappa$ plateau (4 networks) | | |
| `C650-N05` | 0.00750 | 0.00224 | — |
| `C850-N04` | 0.00631 | 0.00168 | — |
| `C850-N05` | 0.00631 | 0.00168 | — |
| `C950-N01` | 0.00562 | 0.00150 | — |
| | Finite-$\kappa$ minimum (9 networks) | | |
| `C650-N03` | 0.0607 | 0.0486 | 0.315 |
| `C650-N04` | 0.103 | 0.0955 | 1.14 |
| `C750-N02` | 0.0756 | 0.0661 | 0.687 |
| `C750-N04` | 0.0847 | 0.0765 | 0.922 |
| `C850-N01` | 0.0781 | 0.0701 | 0.978 |
| `C850-N03` | 0.0462 | 0.0351 | 0.252 |
| `C950-N03` | 0.0701 | 0.0620 | 0.927 |
| `C950-N04` | 0.0177 | 0.00770 | 0.0041 |
| `C950-N05` | 0.0694 | 0.0612 | 0.898 |
| | Large-$\kappa$ plateau (7 networks) | | |
| `C650-N01` | 0.122 | 0.114 | 2.03 |
| `C650-N02` | 0.122 | 0.114 | 2.03 |
| `C750-N01` | 0.105 | 0.0967 | 2.00 |
| `C750-N03` | 0.105 | 0.0967 | 2.00 |
| `C750-N05` | 0.105 | 0.0967 | 2.00 |
| `C850-N02` | 0.0924 | 0.0842 | 1.98 |
| `C950-N02` | 0.0825 | 0.0746 | 1.97 |
